# Commonly used compositional data analysis implementations are not advantageous in microbial differential abundance analyses benchmarked against biological ground truth

**DOI:** 10.1101/2025.02.13.638109

**Authors:** Samuel D. Gamboa-Tuz, Marcel Ramos, Eric Franzosa, Curtis Huttenhower, Nicola Segata, Sehyun Oh, Levi Waldron

## Abstract

Previous benchmarking of differential abundance (DA) analysis methods in microbiome studies have employed synthetic data, simulations, and “real data” examples, but to the best of our knowledge, none have yet employed experimental data with known “ground truth” differential abundance. A key debate in the field centers on whether compositional methods are necessary for DA analysis, which is challenging to answer due to the lack of ground truth data. To address this gap, we created the Bioconductor data package *MicrobiomeBenchmarkData*, featuring three microbiome datasets with established biological ground truths: 1) diverse oral microbiomes from supragingival and subgingival plaques, expected to favor aerobic and anaerobic bacteria, respectively, 2) low-diversity microbiomes from healthy vaginas and bacterial vaginosis, conditions that have been well-characterized through cell culture and microscopy, and 3) a spike-in dataset with constant, known absolute abundances of three bacteria. We benchmarked 17 DA approaches and demonstrated that compositional DA methods are not beneficial but rather lack sensitivity, show increased variability in constant-abundance spike-ins, and, most surprisingly, more frequently produce paradoxical results with DA in the wrong direction for the low-diversity microbiome. Conversely, commonly used methods in microbiome literature, such as *LEfSe*, the Wilcoxon test, and RNA-seq-derived methods, performed best. We conclude that researchers continue using widely adopted non-parametric or RNA-seq DA methods and that further development of compositional methods includes benchmarking against datasets with known biological ground truth.

## Introduction

Compositional methods for microbiome differential abundance (DA) analysis have been developed to address challenges posed by the compositional nature of microbiome data, including sparsity, high dimensionality, and technical zeros (Lin and Peddada, 2020b; Brill *et al*., 2022). Whereas for supervised learning methods, benchmarking against biological “ground truth” can be done through cross-validation, biological ground truth is much more difficult to establish for benchmarking of statistical inference (Friedrich and Friede, 2024). In the context of microbiome DA, several authors have argued that traditional statistical approaches such as linear regression and non-parametric tests should be avoided because of the compositional nature of microbiome data(Tsilimigras and Fodor, 2016; Gloor and Reid, 2016; Friedrich and Friede, 2024; Gloor *et al*., 2016, 2017). However, arguments for using compositional analysis methods for microbiome data have relied on theoretical arguments, simulations, and toy examples. Conversely, elementary statistical methods based on linear regression and non-parametric tests have been reported to show higher consistency and replicability across multiple studies without compromising sensitivity, relative to microbiome-specific and compositional methods (Pelto *et al*., 2024). Other benchmarking efforts have reported a range of results when benchmarking compositional and non-compositional methods: good reproducibility properties of compositional methods but recommended comparison of results from multiple methods (Nearing *et al*., 2022); that only classical statistical methods (linear models, the Wilcoxon test, t-test), limma, and fastANCOM properly control false discoveries at relatively high sensitivity(Wirbel et al.,2024); that certain methods consistently result in higher concordances(Wallen, 2021) (e.g. ANCOM-BC, LEfSe), and that metagenomeSeq (a microbiome-specific method) and edgeR (an RNA-seq method) exhibit low false positive rates and have good power at differing levels of sparsity in in-silico, spike-in, and permuted datasets(Thorsen *et al*., 2016). These studies present contradictory findings, but none employed data with known biological ground truth for true positives or true negatives.

Benchmarking studies can be classified into three categories: mock, simulated, and biological(Bokulich *et al*., 2020). Mock and simulated datasets offer the advantage of known outcomes and are ideal for testing software accuracy but often lack the complexity of actual biological samples. However, they can also embed assumptions that favor methods based on the same assumptions. While biological datasets are more representative of real-world conditions, they often lack known ground truth for quantitative benchmarking.

We therefore developed the *MicrobiomeBenchmarkData* data package to provide biological datasets with known ground truth for benchmarking of DA methods. *MicrobiomeBenchmarkData* provides three experimental datasets of varying ecological complexity, each with a ground truth established through experimental methods other than sequencing. We benchmarked the biological performance of common statistical tests and data transformation methods for microbiome DA. We classify DA methods as classical non-parametric methods, compositional analysis, methods explicitly developed for microbiome data, and approaches borrowed from RNA-seq and single-cell RNA-seq data analysis. We evaluated the performance of each method based on the numbers of true positives and false positives and the variance of spike-in bacteria, and found that compositional and non-compositional methods perform similarly in high-complexity oral microbiomes, but produce inferior and paradoxical results in low-complexity vaginal microbiota, and that centered log-ratio transformation increased variance of constant-abundance spike-in samples. We conclude with a discussion of how microbiome sequencing data violate assumptions of classical compositional data analysis methods, and remind readers of the importance of considering biological relevance in the benchmarking of statistical methods.

## Results

### Microbiome datasets with ground truth

The *MicrobiomeBenchmarkData* package provides three distinct biological scenarios for benchmarking (**Table 1**): 1) subgingival vs. supragingival dental plaque, where the oxygen-poor environment of subgingival plaque favors anaerobic bacteria while the more oxygenated environment of supragingival plaque favors aerobic bacteria (Thurnheer *et al*., 2016), 2) healthy vaginal microbiome vs. bacterial vaginosis, an overgrowth of diverse bacteria in a normally *Lactobacillus*-dominated environment, and 3) spike-in bacteria in stem cell transplantation patient samples(Stämmler *et al*., 2016). Each dataset includes abundance counts, sample metadata, microbial taxonomy, and established ground truth, available as *TreeSummarizedExperiment* objects in R.

**Table 1.**
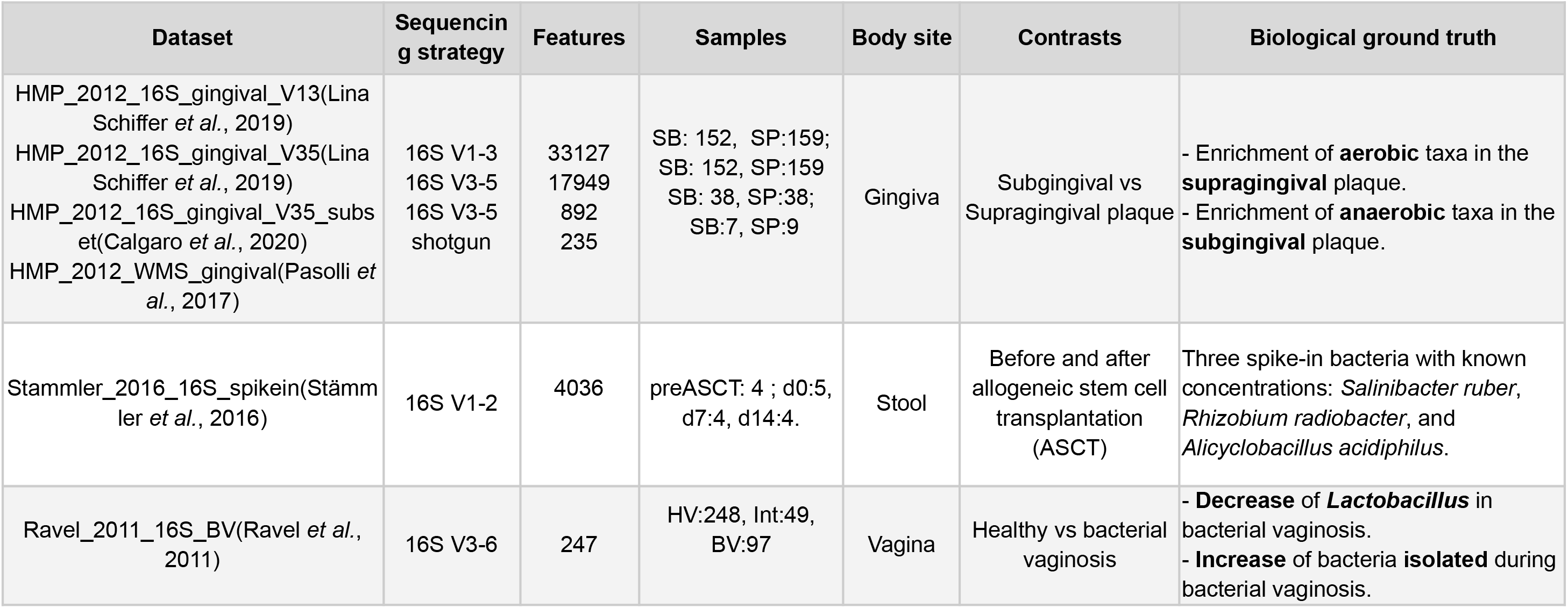
Datasets with biological ground truth in the *MicrobiomeBenchmarkData* package.

#### High complexity dataset: subgingival and supragingival plaque

We utilized Human Microbiome Project (HMP1) data (Human Microbiome Project Consortium, 2012) comprising both 16S rRNA and whole metagenomic shotgun sequencing (WMS) from dental plaque samples. There are two measurements for most participants, sampled in one visit, one from the supragingival and one from the subgingival oral plaque, and some unpaired observations. The biological ground truth stems from differential oxygen availability: supragingival plaques favor aerobic bacteria, while subgingival plaques favor anaerobic bacteria(Calgaro *et al*., 2020; Thurnheer *et al*., 2016). While there is no exact list of differentially abundant taxa, high-performing DA methods can be expected to identify aerobic taxa preferentially abundant in the supragingival plaque and anaerobic taxa preferentially abundant in the subgingival plaque. MicrobiomeBenchmarkData provides annotations of the samples and of oxygen utilization at the genus level, as previously reported(Beghini *et al*., 2019). A naive or random method would find anaerobic and aerobic taxa equally likely to be increased in the supragingival or subgingival plaque.

#### Low-complexity dataset: bacterial vaginosis

The microbiome of the healthy vagina (HV) is characterized by low diversity and a high abundance of species of the *Lactobacillus* genus (Ravel *et al*., 2011, 2013), while Bacterial Vaginosis (BV) is associated with increased vaginal microbial diversity, with BV-affected communities showing nearly three times more OTUs than healthy vaginal communities(Oakley *et al*., 2008). Importantly, HV and BV-associated genera have been established by culture and microscopy, without use of sequencing. This dataset includes Nugent scores (Nugent *et al*., 1991), a Gram stain microscopy-based scoring system for BV classification, for 16S profiles from 248 HV samples, 97 BV samples, and 49 intermediate samples. The established ground truth is the decreased *Lactobacillus* abundance and increased BV-associated genera in BV samples. MicrobiomeBenchmarkData annotates BV-associated bacteria, including *Gardnerella* and *Prevotella*, that have been identified through culture-based experiments (**Supplementary Table 1**) (Alves *et al*., 2014; Patterson *et al*., 2010; Łaniewski and Herbst-Kralovetz, 2021; Bordigoni *et al*., 2020; Onderdonk *et al*., 2016).

#### Known concentration: spike-in bacteria

This dataset comprises microbiome samples from five allogeneic stem cell transplantation (ASCT) patients collected at multiple timepoints. Before library preparation and 16S sequencing, three non-human bacterial species were added in constant absolute amounts to each sample: *Salinibacter ruber, Rhizobium radiobacter*, and *Alicyclobacillus acidiphilus*. This dataset is intended for benchmarking normalization methods such as relative abundance and centered log-ratio: a desirable property of normalization would be to minimize the observed variability of these taxa relative to other taxa that are naturally present in the samples in non-constant absolute abundance.

### Benchmarking DA methods

We evaluated seventeen DA approaches, which are combinations of eleven statistical tests/methods, five normalization/transformation techniques, and three weighting options.

These approaches are classified into five categories based on their primary purpose and target data types (**Supplementary Table 2**).

1. Two classical statistical tests (Classical): Wilcox + TSS and LEfSe +TSS (Segata *et al*., 2011; Khleborodova *et al*., 2022).
2. Five compositional data methods (Compositional): Wilcox + CLR, LEfSe + CLR, ZINQ + CLR (Ling *et al*., 2021), ALDEx2(Fernandes *et al*., 2013, 2014), and ANCOM-BC(Mandal *et al*., 2015; Lin and Peddada, 2020a).
3. Two microbiome-specific methods (Micro): MetagenomeSeq + CSS (Paulson *et al*., 2013) and ZINQ + TSS (Ling *et al*., 2021).
4. Three RNA-seq methods (RNAseq): DESeq2 + Poscounts(Love *et al*., 2014), edgeR + TMM (Robinson *et al*., 2010; McCarthy *et al*., 2012), and limma-voom + TMM (Law *et al*., 2014).
5. Five single-cell RNA-seq methods (scRNAseq): edgeR + TMM + Zimbwave, DESeq2 + Poscounts + Zimbwave, limma-voom + TMM + Zimbwave (Risso *et al*., 2018; Van den Berge *et al*., 2018; Calgaro *et al*., 2020), MAST(McDavid *et al*., 2022; Finak *et al*., 2015), and Seurat (Hao *et al*., 2021; Stuart *et al*., 2019; Butler *et al*., 2018; Satija *et al*., 2015).

#### Benchmark 1: Dental plaque microbiome analysis

Using matched subgingival and supragingival plaque samples, we identified DA taxa (FDR < 0.1) (**Figure 1**). All methods effectively identified strong enrichment of anaerobic bacteria in subgingival plaque (**Figure 1a**, bottom panel). However, only seven methods (Wilcox+TSS, LEfSe+TSS, LEfSe+CLR, ZINQ+TSS, DESeq2+Poscounts, limma-voom+TMM, and MAST) detected a significant enrichment of aerobic bacteria in supragingival plaque vs subgingival plaque (p < 0.05, Fisher’s Exact Test) (**Figure 1a**, top panel). The classical non-parametric Wilcoxon test, and closely related LEfSe, detected the most aerobic taxa as DA in supragingival plaque, but also identified greater numbers of facultative and obligate anaerobes. When evaluating across a range of thresholds for calling a taxon DA, calling between 1 and 50 taxa DA (**Figure 1b**), differences between the methods were minor.

**Figure 1.**
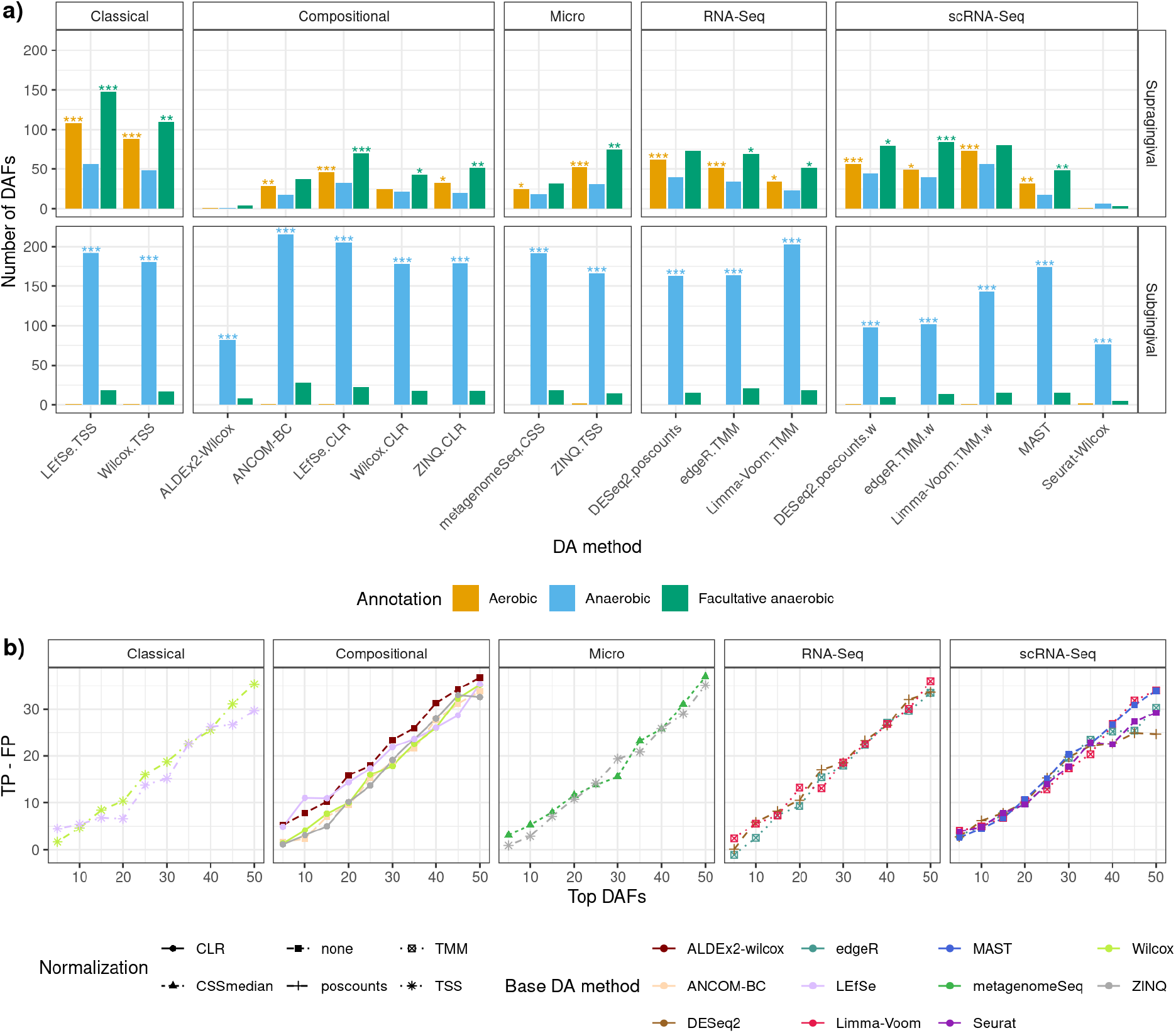
Benchmark DA approaches using subgingival versus supragingival plaque microbiomes. Seventeen differential abundance (DA) approaches were applied to the dental plaque dataset (*HMP_2012_16S_gingival_V35*). **a)** Enrichment pattern of differentially abundant features (DAFs) (FDR ≤ 0.1) classified as aerobic (mustard), anaerobic (blue), or facultative anaerobic (green) in supragingival (upper panel) and subgingival (lower panel) samples. Significance levels from enrichment testing (hypergeometric test) are indicated by asterisks: *P ≤ 0.05, **P ≤ 0.01, ***P ≤ 0.001. **b)** Performance assessment (the difference between putative TPs and FPs) is displayed for top DAFs (ranked by effect size, independent of statistical significance), grouped by base method and transformation approach.

#### Benchmark 2: Bacterial vaginosis analysis

All methods detected BV-associated taxa in BV samples (**Figure 2a**), though the statistical significance of the enrichment analysis was limited by sample complexity. Most methods, except ALDEx2 and Seurat, correctly identified *Lactobacillus* enrichment in HV samples. All compositional methods incorrectly identified one or more BV-associated taxa as over-abundant in healthy samples, while among non-compositional methods only DESeq2 with the poscounts normalization, and metagenomeSeq, made the same paradoxical error. ALDEx2 (a compositional method) was the only method to paradoxically identify *Lactobacillus*, the dominant genus in healthy vaginal microbiomes, as a marker of BV (**Figure 2a**). Overall, LEfSE and Wilcoxon test with TSS normalization, and edgeR with TMM (weighted or unweighted) showed the highest overall sensitivity without any paradoxical errors. These overall conclusions are consistent in threshold-free comparisons of True Positives minus False Positives from 0 to 20 top DA taxa (**Figure 2b**). It is worth noting that while LEfSE uses a p-value cutoff from the Wilcoxon test, it ranks by LDA coefficients instead of by p-values as when using the Wilcoxon test on its own.

**Figure 2.**
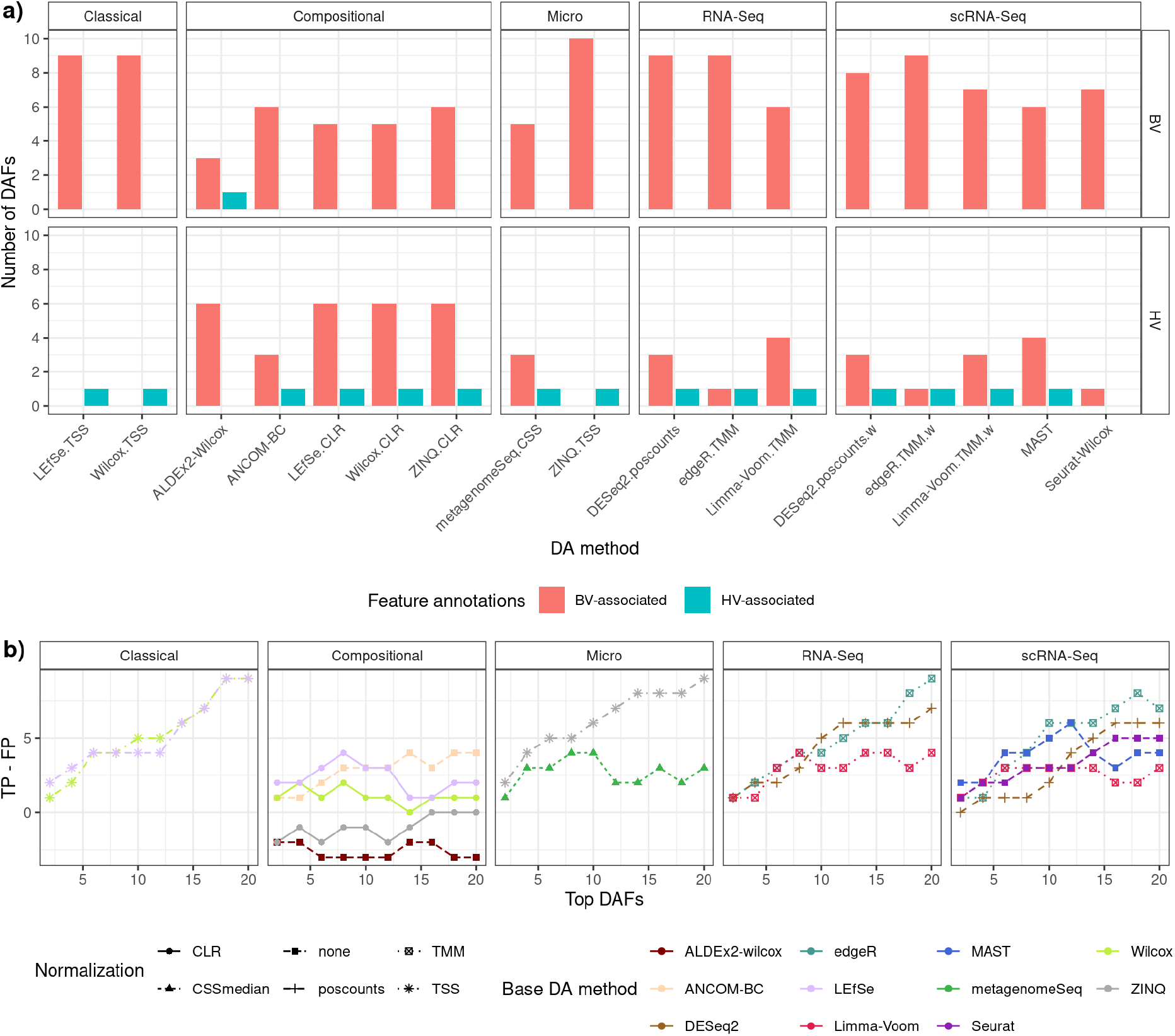
Benchmark DA approaches using healthy vaginal versus bacterial vaginosis microbiomes. Seventeen differential abundance (DA) approaches were applied to the dataset with vaginal microbiome samples (*Ravel_2011_16S_BV*). **a)** The numbers of DA taxa (FDR ≤ 0.1) annotated as HV-associated (blue) or BV-associated (red) from BV (upper panel) and HV (lower panel) samples. No enrichment results achieved statistical significance (hypergeometric test P ≥ 0.05). **b)** The accuracy (putative TP-FP) for top DA taxa, comparing performance across base method and transformation approach.

To clarify the paradoxical differential abundance observed with CLR normalization in low-complexity vaginal samples, we plotted TSS and CLR-transformed abundances of seven taxa whose differential abundances (DA) changed with CLR normalization (**Figure 3**). The over-abundance of bacterial vaginosis (BV)-associated genera in healthy vagina samples is evident in the CLR-transformed data but not in log(counts + 1) or log(TSS + 1) data. This effect is further illustrated by scatter plots comparing TSS and CLR values for two non-paradoxical taxa (*Lactobacillus* and *Prevotella)* and two paradoxical taxa (*Streptococcus* and *Corynebacterium*) (**Figure 4**). For the high-abundance *Lactobacillus* and *Prevotella* genera which did not demonstrate paradoxical effects, there is a strong positive correlation between TSS and CLR-transformed data. Conversely, for low-abundance BV-associated taxa (*Corynebacterium* and *Streptococcus*), CLR-transformed abundance is primarily influenced by the dominant *Lactobacillus*. This results in paradoxically high CLR-transformed abundance of these taxa in healthy samples compared to those with BV. The high relative abundance of *Lactobacillus* in healthy vaginas amplifies the CLR-transformed abundances of *Streptococcus* and *Corynebacterium*, leading to the misleading appearance of increased BV-associated taxa in healthy samples (**Supplementary Table 3**).

**Figure 3.**
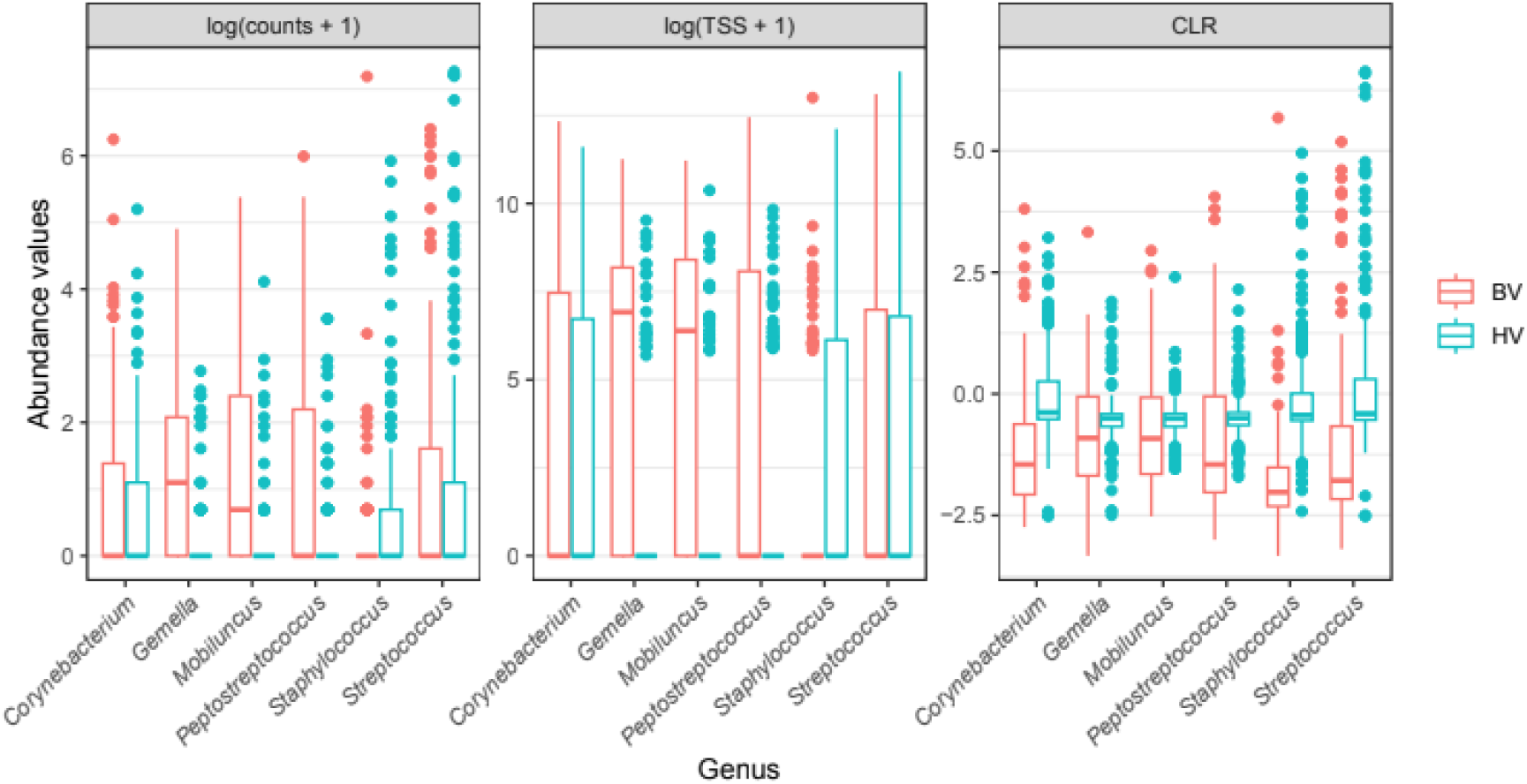
The impact of CLR transformation on the abundance patterns of low-abundant genera in low-complexity microbiome data. Comparison of the BV-associated taxa abundance between HV (blue) and BV (red) samples under three normalization approaches: log-transformed raw counts (left), TSS (center), and CLR (right). Log-transformed abundance values (y-axis) show how CLR transformation flips the abundance values for low-abundant, BV-associated genera.

**Figure 4.**
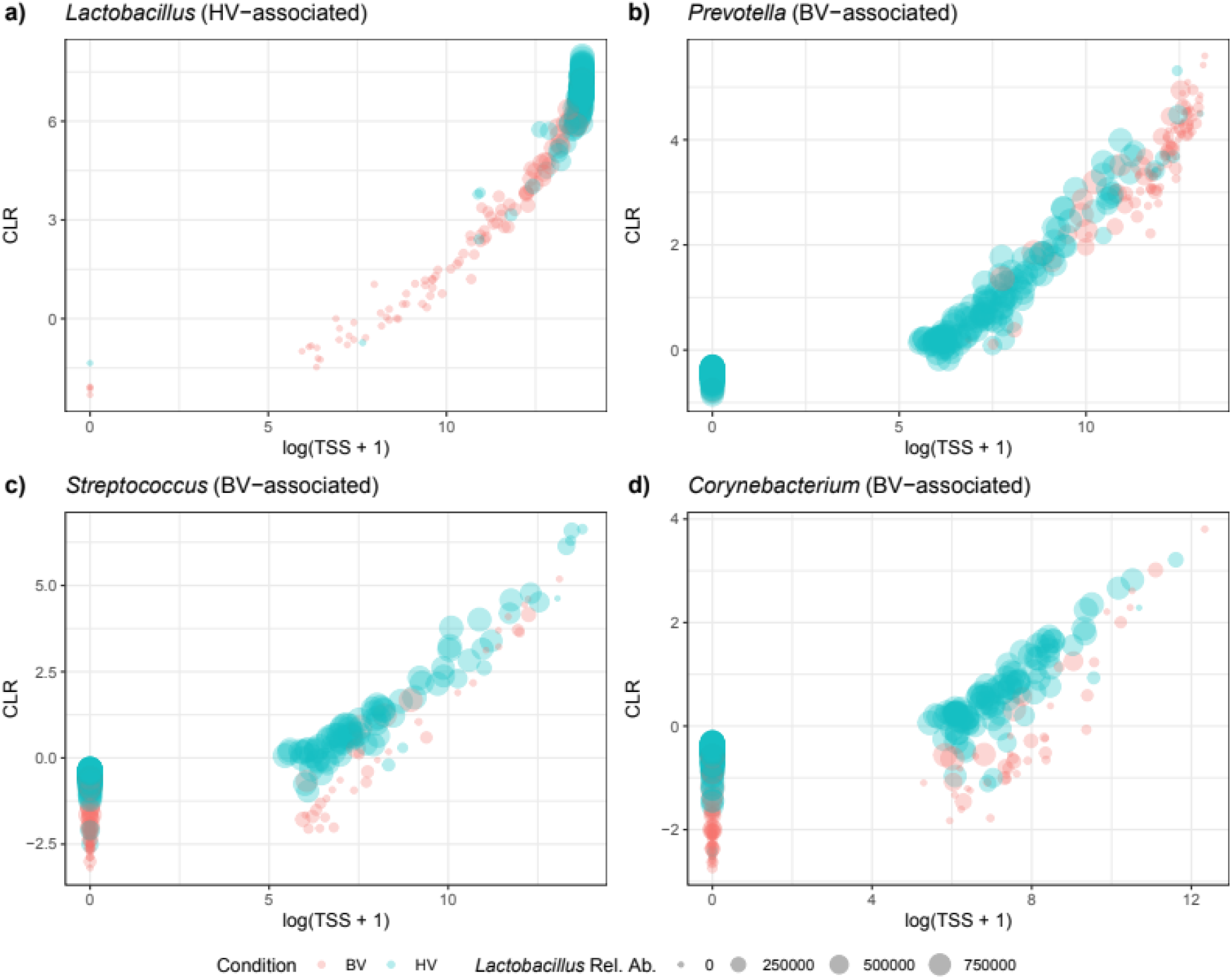
CLR transformation inflates the abundance of BV-associated genera in HV samples. Comparison of TSS versus CLR transformed abundances for high and low abundance taxa. Data points represent individual taxa (indicated in plot titles), with the size of points proportional to the relative abundance of *Lactobacillus*. Abundant HV and BV-associated taxa (*Lactobacillus* and *Prevotella*, respectively) show a linear relationship between log(TSS+1) and CLR, agreeing with the biological ground truth; *Lactobacillus* is more abundant in HV (panel a) and *Prevotella* is more abundant in BV (panel b). The two BV-associated taxa (panels c and d) are similarly low-abundant in both HV and BV samples (i.e., the majority of samples below log(TSS+1) <8); however, CLR-transformation disproportionally amplifies the abundance value of them depending on the abundance of *Lactobacillus*, making BV-associated taxa look more abundant in HV samples (large blue circles) than BV-samples (small red circles).

#### Benchmark 3: Spike-in analysis

We used the spike-in dataset to benchmark relative abundance against compositional normalization for observed stability of the taxa with constant absolute abundance vs. naturally present taxa of varying abundance. We compared TSS and GMN (Geometric Mean Normalization) because both methods avoid log transformation, allowing for more comparable Coefficient of Variation (CV) values. TSS yielded lower CV values for bacteria maintained at constant absolute abundance than GMN, in both absolute CV and rank among all taxa (**Table 2**). This indicates that compositional data analysis methods, like CLR transformation, may introduce greater measurement instability than relative abundance methods. Furthermore, when the features were ranked based on their CV values obtained with each normalization method, all three spike-in bacteria showed lower ranks when using TSS normalization (**Table 2 and Figure 5**). which indicates less variation using the TSS method.

**Table 2.** Variation of spiked-in bacteria with different pre-processing methods. Two preprocessing methods, Total Sum Scale (TSS) and Geometric mean (GM), were applied to the Stammler_2016_16S_spikein dataset. The coefficient of variation (CV) and standard deviation (SE) of the abundance of the three spike-in bacteria were calculated.

| Species | Preprocessing | CV Ranking | CV | SE |
| --- | --- | --- | --- | --- |
| *A. acidiphilus* | TSS | 2836 | 0.62 | 0.09 |
|  | GMN | 3076 | 0.74 | 0.11 |
| *R. radiobacter* | TSS | 2945 | 0.7 | 0.09 |
|  | GMN | 3129 | 0.82 | 0.11 |
| *S. ruber* | TSS | 2901 | 0.65 | 0.09 |
|  | GMN | 3030 | 0.71 | 0.1 |

**Figure 5.**
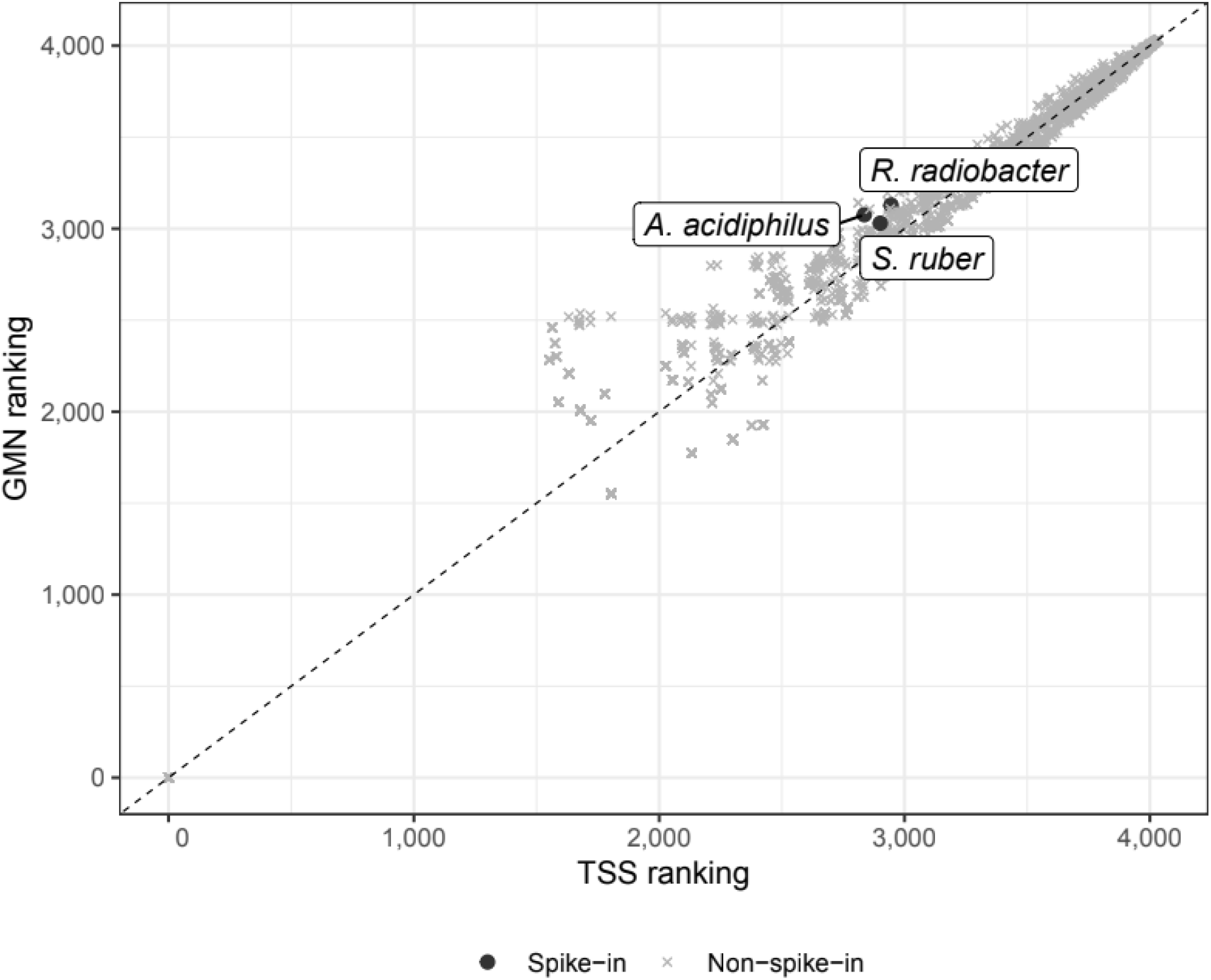
Comparison of bacterial rankings in the spike-in dataset based on their coefficient of variation across all samples. **A)** Dots represent features in the spike-in dataset (*Stammler_2016_16S_spikein*).

## Discussion

The *MicrobiomeBenchmarkData* R/Bioconductor package provides three biologically distinct microbiome datasets with lab-verified biological ground truth, enabling biologically-grounded benchmarking of DA methods. We use these datasets to address a key question in the field: **do microbiome data require the use of compositional data analysis (CoDA) methods?**

Microbiome data share important characteristics with classical compositional data, representing relative rather than absolute abundances(Odintsova *et al*., 2017). As a result, CoDA methods, such as log-ratio transformation (LRT), are presumed by many to be superior to non-compositional normalization methods. However, microbiome sequencing differs from classical compositional data in several ways: the true number of taxa is unknown, there are varying proportions of unannotated reads, and library size variations challenge strict adherence to compositionality(Jeganathan and Holmes, 2021; Greenacre *et al*., 2023). Additionally, microbiome sequencing data presents significant challenges for CoDA, including sparsity, zero handling(Jeganathan and Holmes, 2021; Greenacre *et al*., 2023), and interpretation difficulties in high-dimensional data(Greenacre *et al*., 2023). Some studies have also highlighted poor performance of CoDA methods(Calgaro *et al*., 2020) and advocated for other classical statistical approaches (Love *et al*., 2014; Robinson *et al*., 2010; McMurdie and Holmes, 2014). Overall, our benchmarking in datasets with biological ground truth finds that, contrary to widely-held beliefs, traditional non-compositional approaches may better identify biologically relevant features, *particularly* in low-complexity microbiomes.

### Effects of microbial community complexity on DA analysis

Compositional effects are less pronounced in high-complexity ecosystems without dominant species. It is therefore unsurprising that in complex oral microbiome samples, differences between compositional and non-compositional methods were minor in threshold-free comparisons, with differences mainly in the sensitivity-specificity / precision-recall tradeoff. More surprisingly, in low-complexity vaginal samples where *Lactobacillus* is usually dominant, we observed poor performance and paradoxical results from the compositional methods tested. While all methods except for ALDEx2-Wilcox correctly detected higher *Lactobacillus* abundance in healthy samples, compositional methods frequently identified BV-associated taxa as also having increased abundance in healthy samples. This resulted from CLR transformation artificially inflating the apparent abundance of low relative-abundance BV-associated genera in healthy samples where *Lactobacillus* dominates. We even observed the direction reversals in CLR-transformed BV markers *Streptococcus* and *Corynebacterium*, demonstrating that compositional transformation can produce paradoxical results in low-diversity microbial datasets. Geometric mean transformation, intended to correct for compositional effects, also artificially increased variability in three spike-in bacteria across samples compared to TSS.

### Limitations

The *MicrobiomeBenchmarkData* package and these analyses could be strengthened by expansion to more than three representative datasets and to include whole metagenomic shotgun sequencing data. The current analysis challenges assumptions that CoDA methods are superior for microbiome data but should be expanded in the future to confirm and add detail to these conclusions.

## Conclusions

We benchmarked DA analysis methods using microbiome datasets with ground truths provided through the *MicrobiomeBenchmarkData* data package and gained three key insights. First, community complexity strongly influences method performance - while RNA-seq-derived methods excel in analyzing complex, high-diversity communities, no single method performs optimally across all scenarios. Second, compositional methods, particularly CLR transformation, can produce misleading results in low-diversity communities by artificially inflating the apparent abundance of low relative-abundance taxa. Third, simpler normalization approaches like TSS can introduce less error than centered log-ratio transformation, especially when analyzing low-abundant taxa in low-complexity communities. These results highlight the need for biologically grounded benchmarking in selecting analysis methods, and do not support the use of CLR or other commonly-used implementations of compositional data analysis for microbiome community sequencing data.

## Materials and Methods

### Datasets

The *MicrobiomeBenchmarkData* package provides three biologically distinct datasets. First, we obtained gingiva data from the HMP1 project(Human Microbiome Project Consortium, 2012) through three sources:

1. Complete 16S data (*HMP_2012_16S_gingiva_V13* and *HMP_2012_16S_gingiva_V35* datasets) from the *HMP16SData* package (ver.1)(Lucas Schiffer *et al*., 2019)
2. A subset of the HMP1 16S V3-5 data (*HMP_2012_16S_gingiva_V35_subset*) previously used in a benchmarking study(Calgaro *et al*., 2020), accessed via GitHub(Calgaro)
3. Whole metagenomic shotgun sequencing (WMS) data (*HMP_2012_WMS_gingival*) from ten subjects, obtained through the *curatedMetagenomicData* package (ver.3)(Pasolli *et al*., 2017)

The bacterial vaginosis dataset (*Ravel_2011_16S_BV*) is a cross-sectional study of reproductive-age women(Ravel *et al*., 2011). A spike-in bacteria dataset from ASCT patients(Stämmler *et al*., 2016), contains three bacterial species (*S. ruber, R. radiobacter*, and *A. acidiphilus*) at known concentrations (equivalent to 3.0 x10^8^, 5.0×10^8^, and 1.0×10^8^ 16S rDNA copies for each species, respectively). Both bacterial vaginosis and spike-in bacteria data were obtained from the supplementary materials provided by the original literature.

Vaginal samples were classified as ‘healthy’, ‘intermediate’, and ‘bacterial vaginosis’ based on whether their reported Nugent scores were low (0-3), medium (4-6), or high (7-10), respectively.

### Differential abundance analysis

#### Data annotation and processing

Taxa from the gingival plaques dataset were annotated at the genus level following a previous report(Beghini *et al*., 2019). Samples in the *Ravel_2011_16S_BV* were labeled as healthy, intermediate, and bacterial vaginosis based on their Nugent scores: 0-3 (low), 4-6 (intermediate), and 7-10 (high), respectively.

*Ravel_2011_16S_BV* counts data were summarized at the genus level. We filtered out taxa with less than 20% prevalence (a count of at least one was considered as a presence) for both the *HMP_2012_16S_gingival_V35* and the *Ravel_2011_16S_BV* datasets.

#### Normalization methods

We implemented the normalization techniques most commonly used for the target methods: Total Sum Scaling (TSS; also referred to as relative abundance) divides each feature’s count by the total sum of counts in the sample and multiplied by 10^6^, converting counts to relative proportions. Cumulative sum scaling (CSS), proposed for microbiome data(Paulson *et al*., 2013), adjusts for sequencing depth differences by calculating sample-specific scaling factors based on the cumulative sum of ordered relative abundances up to a data-driven determined quantile, aiming to reduce non-biological between-sample variations and improve comparability between samples. Additive Log-Ratio (ALR) transformation, proposed for compositional data (Mandal *et al*., 2015; Fernandes *et al*., 2014), converts relative abundances to log-ratios by selecting a reference taxon and calculating the logarithm of the ratio of each component to this reference, allowing for the application of standard statistical methods to compositional data. Centered log-ratio (CLR) transformation, also proposed for compositional data analysis (Aitchison, 1982), normalizes data by dividing each feature’s abundance by the geometric mean of all features in the sample, followed by log transformation. Trimmed mean of M-values (TMM) (Robinson and Oshlack, 2010) normalization estimates relative abundance production levels between samples by calculating a scaling factor based on the trimmed mean of log-fold changes, assuming most features are not differentially expressed. Relative log expression (RLE) normalization (Love *et al*., 2014) uses the median ratio of each feature’s expression to the geometric mean across all samples as a scaling factor.

#### Statistical analysis

We applied seventeen combinations of DA and normalization methods (**Supplementary Table 2**) to the dental plaque and BV datasets using the *benchdamic* package (version 1.3.0)(Calgaro *et al*., 2022). The significant threshold for DA features was set at FDR ≤ 0.1. We calculated the effect size using a log-fold change of the normalized counts for the Wilcoxon test, LEfSe (implemented as the *lefser* package (version 1.14.0)), and ZINQ. For other approaches, we used method-specific calculations.

#### Enrichment analysis

We performed bug set enrichment analysis on gingival plaque and vaginal datasets. We used the hypergeometric test from the *benchdamic* package (version 1.3.0)(Calgaro *et al*., 2022, 2020) with a significance threshold of p < 0.05.

### Performance Evaluation

#### Putative True/False Positive analysis

Features were ranked by effect size and evaluated across various threshold cutoffs. We defined the putative true/false positives as

**Putative true positives**:

- Gingival dataset: over-abundant aerobic taxa in supragingival plaque **plus** over-abundant anaerobic taxa in subgingival plaque
- Vaginal dataset: over-abundant HV-associated taxa in HV samples **plus** over-abundant BV-associated taxa in the BV samples

**Putative false positives**:

- Gingival dataset: over-abundant aerobic taxa in subgingival plaque **plus** over-abundant anaerobic taxa in supragingival plaque
- Vaginal dataset: over-abundant HV-associated taxa in BV samples **plus** over-abundant HV-associated taxa in the BV samples

Analysis of the difference between putative true positives and putative false positives (TP - FP) was carried out as described in (Calgaro *et al*., 2020) using the *benchdamic* package (version 1.3.0). After running the DA methods, the features were ranked based on adjusted p-value (FDR), except for the features analyzed with the ‘LEfSe + TSS’ and ‘LEfSe + CLR’ methods, which were ranked based on LDA scores. This ranking was performed without applying any threshold of statistical power.

#### Coefficient of variation

The count data in the *Stammler_2016_16S_spikein* dataset were normalized using Total Sum Scaling (TSS; Equation 1) and geometric mean normalization (GMN; Equation 2). Coefficients of variation (CV) and standard errors (SE) were calculated from 1,000 non-parametric bootstrap replicates with replacement for each feature, considering all samples irrespective of treatment or time.

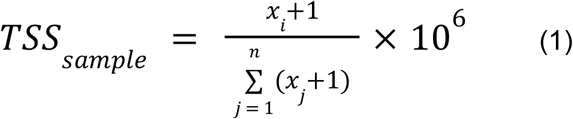

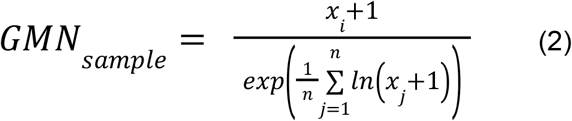

## Data and code availability

- Data as text files: available on Zenodo https://doi.org/10.5281/zenodo.6911026
- Data with additional functionality as *MicrobiomeBenchmarkData* R package: available through Bioconductor https://www.bioconductor.org/packages/MicrobiomeBenchmarkData/
- Dataset preparation scripts: https://github.com/waldronlab/MicrobiomeBenchmarkDataPrep
- Code to reproduce analyses in this manuscript: https://waldronlab.io/MicrobiomeBenchmarkDataAnalyses

## Supplementary Figures

**Supplementary Figure S1.**
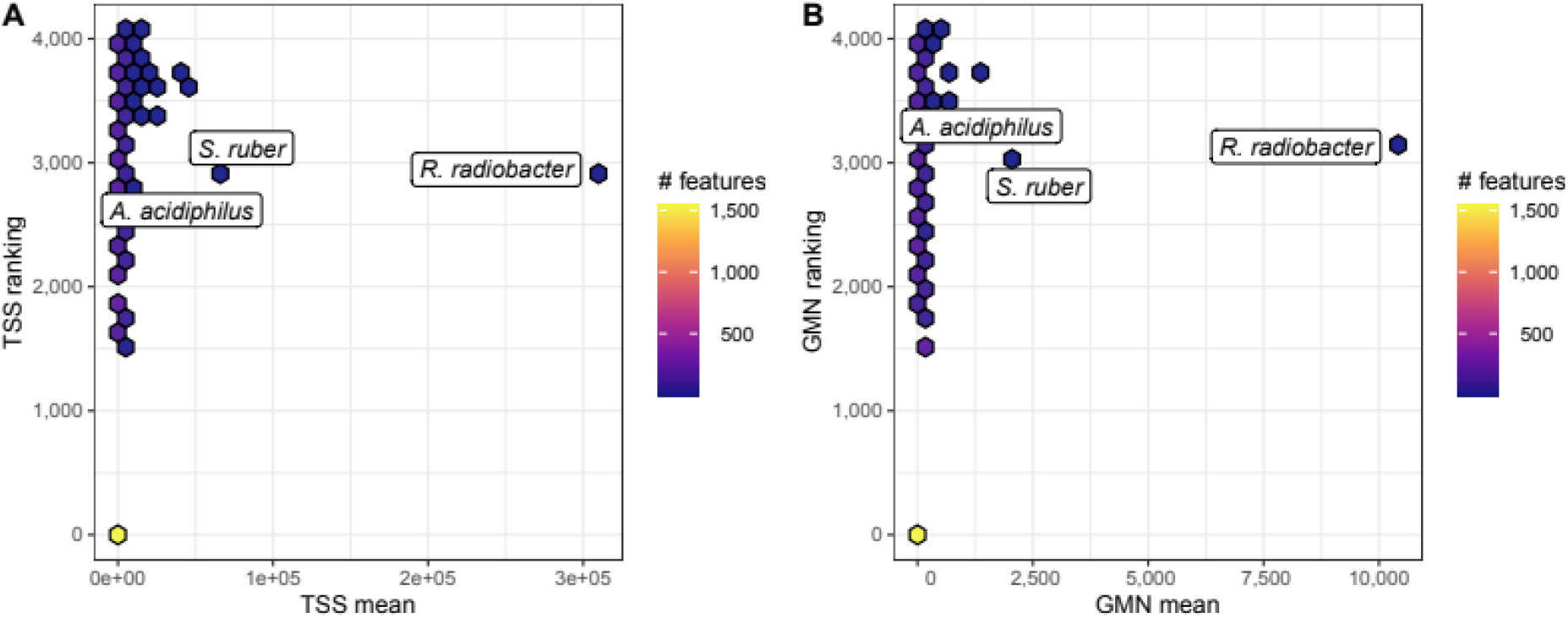
Comparison of CV and mean relative abundance (TSS/GM) versus ranking (Stammler_2016_16S_spikein). Counts were normalized with either **a)** Total Sum Scaling (TSS) or **b)** by dividing by the geometric mean (geometric mean normalization; GMN). The coefficient of variation was estimated for each taxa and they were ranked bsed on the results.

## Notes

https://www.bioconductor.org/packages/MicrobiomeBenchmarkData/

### Competing Interest Statement

The authors have declared no competing interest.

